# The relationship between brain activation and mitochondrial complex I protein levels during cognitive function in healthy humans: an [18F]BCPP-EF PET and functional MRI study of task switching

**DOI:** 10.1101/2024.09.25.614887

**Authors:** Ekaterina Shatalina, Thomas Whitehurst, Ellis Chika Onwordi, Alexander Whittington, Ayla Mansur, Atheeshaan Arumuham, Tiago Reis Marques, Roger N. Gunn, Sridhar Natesan, Matthew M. Nour, Eugenii A. Rabiner, Matthew B. Wall, Oliver D Howes

## Abstract

Mitochondrial complex I is the largest enzyme complex in the respiratory chain and can be non-invasively measured using [18F]BCPP-EF positron emission tomography (PET). Neurological conditions associated with mitochondria complex I pathology are also associated with altered blood oxygen level dependent (BOLD) response and impairments in cognition. To evaluate the link between mitochondrial complex I, cognition and associated neural activity, 23 cognitively healthy adults underwent a [18F]BCPP-EF PET scan and a functional magnetic resonance imaging (fMRI) scan during which they performed a task switching exercise.

We found significant positive associations between [18F]BCPP-EF volume of distribution (VT), which measures mitochondrial complex I levels and the task switching fMRI response (Partial Least Squares (PLS) Canonical Analysis (CA), first component r=0.51, p=0.03). Exploratory Pearson’s correlations showed significant positive associations between mitochondrial complex I levels and the fMRI response in regions including the dorsolateral prefrontal cortex (*r*=0.61, *p*=0.0019), insula (*r*=0.46, *p*=0.0264) parietal-precuneus (*r*=0.51, *p*=0.0139) and anterior cingulate cortex (*r*=0.45, *p*=0.0293). Mitochondrial complex I levels across task-relevant regions were also predictive of task switching accuracy (PLS-Regression (PLS-R), R2=0.48, RMSE=0.154, p=0.011) and of switch cost (PLS-R, R^2=0.38, RMSE=0.07, p=0.048).

Our findings suggest that higher mitochondrial complex I levels may underlie an individual’s ability to exhibit a stronger BOLD response during task switching and are predictive of better task switching performance. This provides the first evidence linking the BOLD response with mitochondrial complex I and suggests a possible biological mechanism for aberrant BOLD response in conditions associated with mitochondrial complex I dysfunction, that should be tested in future studies.

## Introduction

Mitochondrial complex I is an electron transport chain super-complex that facilitates the first step of oxidative phosphorylation and plays a key role in neuronal energy production [1–4]. Mitochondrial complex I abnormalities have been implicated in multiple neurological and psychiatric disorders [5–9] that are also associated with aberrant neural function both during rest [10–15] and during cognitive tasks [16–20], as measured by functional Magnetic Resonance Imaging (fMRI).

FMRI relies on imaging the Blood Oxygen Level Dependent (BOLD) signal, which measures the local haemodynamic response to changing oxygen demand driven by changes in neural activity. The BOLD signal change has been shown to reflect the energy demand of synaptic firing and is a key technique that can be used to investigate the relationship between neuronal signalling and measures of metabolism and respiration [21–25]. Specifically, studies employing [13C]-magnetic resonance spectroscopy (MRS) suggest that changes in the BOLD signal during functional activation are closely related to changes in neuronal oxidative metabolism [26]. Furthermore, studies using [18F]Fluorodeoxyglucose Positron Emission Tomography (FDG-PET) show that the BOLD signal has high spatial similarity to glucose metabolism [27–30].

Further evidence comes from pharmacological challenge studies that alter mitochondrial function. For example, pharmacologically inhibiting mitochondrial Ca2+ uniporter (mCU) function impairs calcium uptake into mitochondria which further impairs ATP synthesis [31]. In rats mCU inhibition has been shown to reduce both spontaneous neuronal activity and BOLD signal, measured using intracortical electrodes and fMRI respectively, while pharmacological enhancement of mCU has opposite effects, increasing BOLD signal and spontaneous neuronal activity [32, 33]. Other studies have carried out pharmacological challenges using methylene blue, which enters neural mitochondria and bypasses mitochondrial complex I by accepting electrons from 1,4,-dihydronicotinamide adenine dinucleotide (NADH) and transferring these electrons to mitochondrial complex III [34, 35]. It can thus be used to enhance mitochondrial function and has been investigated as a therapy for cognitive dysfunction in Alzheimer’s disease [36]. In healthy humans, administering a single dose of methylene blue has been shown to increase the BOLD response during psychomotor vigilance and short term memory tasks, and to improve cognitive performance [37]. However, while the evidence above indicates that mitochondrial function influences BOLD measures of neuronal function, it is unclear to what degree natural variation in key mitochondrial components, such as mitochondrial complex I, influences the BOLD signal.

We set out to test the relationship between mitochondrial complex I levels and the BOLD response during fMRI of task switching in healthy adults. Task switching, also known as set shifting, is one of the executive functions of the brain that allows for switching of attention between one task and another [38]. Tasks engaging switching have been shown to reliably activate a large proportion of the brain, including the frontoparietal network [39, 40], which separately has been shown to have the highest metabolic cost, measured by the degree of glucose utilisation, relative to other networks across the brain in a study combining FDG-PET and fMRI [41]. This makes task switching a suitable experimental paradigm for testing the relationship between task fMRI and novel markers of metabolic function. To measure mitochondrial complex I we used novel positron emission tomography tracer, which binds mitochondrial complex I with high affinity. The development of 18F-2-tert-butyl-4-chloro-5- {6-[2-(2-fluoroethoxy)-ethoxy]-pyridin-3-ylmethoxy-2H-pyridazin-3-one ([^18^F]BCPP-EF) has made it possible for the first time to quantify levels of a mitochondrial protein directly in the living human brain [42–48]. We hypothesised that BOLD magnitude in brain regions relevant to task switching will be positively associated with individual differences in mitochondrial complex I levels in those regions, as measured by [18F]BCPP-EF PET [42, 49] across healthy adults.

## Methods

This study was approved by the Administration of Radioactive Substances Advisory Committee (ARSAC, UK) and the London-West London and GTAC ethics committee (Integrated Research Application System reference: 209761, study reference 16/LO/1941) and was carried out in accordance with relevant guidelines and regulations. Participants were recruited through newspaper and local advertisements from within Greater London. All participants received a description of the study before providing written informed consent to participate. Inclusion criteria were: being over 18 and under 65 years-old, capacity to consent to participation in the study, and Allen’s test showing adequate collateral circulation and a normal coagulation test to facilitate arterial blood sampling. In addition to this all participants confirmed they do not have colour-blindness. Exclusion criteria included: history of or current substance use disorder (other than tobacco), history of head injury or neurological abnormality, use of any psychoactive medications, significant physical, psychiatric, or neurological comorbidity, and contraindications to PET or MRI scanning. All subjects had adequate command of English, underwent a structural MRI scan and dynamic PET scan with [^18^F]BCPP-EF, and performed a task switching exercise during an fMRI scan,

### 2.1 Positron emission tomography

[^18^F]BCPP-EF was synthesized as previously described [42]. [18F]BCPP-EF PET scans were conducted at the Invicro Clinical Imaging Centre in London using a Siemens Hi-Rez Biograph 6 PET/CT scanner (Siemens, Erlangen, Germany). Prior to each PET scan, a low-dose CT scan (30 mAs, 130 keV, 0.55 pitch) was carried out for attenuation correction. The radiotracer was administered intravenously as a 20 mL bolus over 20 seconds at the beginning of the scan, via a cannula placed in the cubital or forearm vein. An additional cannula was placed in the radial artery for arterial blood sampling. Dynamic emission data were collected over 90 minutes post-administration and reconstructed into 26 frames (8 frames of 15s, 3 frames of 60s, 5 frames of 120s, 5 frames of 300s, and 5 frames of 600s) using a discrete inverse Fourier transform method, with corrections for attenuation, randoms, and scatter applied.

During the initial 15 minutes of the scan, whole-blood activity was continuously monitored at a rate of 5 mL/min using an automatic blood sampling system (Allogg AB, Mariefred, Sweden). Measurements of total blood and plasma radioactivity were also taken using a Perkin Elmer 1470 10-well gamma-counter (Massachusetts, USA) with samples drawn at the following intervals after post tracer injection: 10, 15, 20, 25, 30, 40, 50, 60, 70, 80, and 90 minutes.

### 2.2 PET Image Analysis and Processing

PET image data was processed using MIAKAT (version 4.3.7), an in-house quantification tool developed by Invicro, which incorporates MATLAB (version R2018b; MathWorks Inc., Massachusetts, USA) and FSL (version 5.0.11; FMRIB) for brain extraction, and SPM12 (Wellcome Trust Centre for Neuroimaging, London, UK) for image segmentation and registration tasks.

The process involved extracting the brain, segmenting grey matter, and performing rigid-body co-registration to standard reference space on each subject’s structural MR image. PET images were then aligned to these MR images and underwent motion correction through frame-to-frame rigid-body registration. Subsequently, each subject’s PET data was transformed into the standard MNI152 space to facilitate kinetic modelling.

### 2.4 Tracer Kinetic Modelling

Time-activity curves (TAC) were fitted in a voxel-wise manner using the multilinear analysis 1 (MA1) model to estimate the total volume of distribution (*V*_T_). Outlier voxels were removed by thresholding images at a *V*_T_ of 55, as in previous work [50], which assumes values above 55 are of supraphysiological level based on previous work using this tracer, no outlier voxels overlapped with preselected ROIs for any subject (details in the supplement and see [51]).

### 2.5 MRI data acquisition

T1-weighted magnetisation-prepared rapid acquisition gradient echo (MPRAGE) images were acquired on a Siemens Magnetom Prisma 3T scanner (Siemens, Erlangen, Germany) using the in-built body coil for Radio Frequency (RF) excitation and the manufacturer’s 64 channel phased-array head/neck coil for reception. These images were used for co-registering the PET and fMRI data. Acquisition parameters were the following: repetition time (TR) = 2300.0 ms, echo time (TE) = 2.28 ms, flip angle = 9°, field of view (FOV) = 256 × 256 mm, 176 sagittal slices of 1 mm thickness, distance factor = 50%, voxel size = 1.0 × 1.0 × 1.0 mm.

Functional data were acquired for a duration of 9 minutes and consisted of T2* weighted transverse echo planar image (EPI) slices. The acquisition consisted of 180 volumes, collected in an interleaved mode in an ascending direction, with GRAPPA (Generalized autocalibrating partially parallel acquisitions) factor = 2 and 2x2x3mm voxel dimensions in the plane, FoV = 250mm, TR = 3000 ms, TE = 30ms, flip angle = 90°, total slices = 44, bandwidth = 1594 Hz/px.

### 2.6 Task switching task design

In the scanner participants completed a switching task adapted from a task previously used in [52]. Participants completed a practice session before the fMRI scan to ensure they understood the rules. During the task pairs of numbers and letters are displayed in either blue or green text (supplementary figure 1). When displayed in green, participants were instructed to focus on the letters and indicate if the letters were vowels or consonants, by pressing a button an MRI-compatible controller as quickly as possible. When the number-letter pairs appeared in blue, the participants were instructed to focus on the numbers, indicating with the controller buttons if they were odd or even.

The task involved two trial types: “switch trials,” where the colour changed from the previous (t-1) trial, and “no-switch trials,” where the colour on the trial (t) was identical to the previous (t-1) trial. Switching trials occurred in the sequence pseudo-randomly. The probability of switch/no-switch on any given trial was uncorrelated to the recent trial history. The task therefore required participants to flexibly switch between two feature-response mappings on each trial, each rendering one stimulus dimension irrelevant, as determined by an abstract rule that required integration of current sensory experience and working memory.

Analysis treated each instance of switching or not as a discrete event, comprising a total of 108 no-switch trials and 42 switch trials. Task performance metrics included switch cost, defined as the difference in average reaction times between switch and no-switch conditions, and switching accuracy, measured as the percentage of correct responses in switch trials [53]. Participants whose switching accuracy fell more than 2.5 standard deviations below the sample mean were excluded from the analysis (*n*=1), assuming disengagement from the task, consistent with criteria used in prior studies [54, 55].

### 2.1 Data preprocessing

FSL (FMRIB Software Library v5.0.4; http://www.fmrib.ox.ac.uk/fsl/) was used for preprocessing of functional data and anatomical data. Brain extraction from anatomical data was carried out using BET, with further preprocessing performed via the fsl_anat script. Motion correction for the functional data was carried out using the FMRIB Linear Image Registration Tool (MCFLIRT), and spatial smoothing was applied using a Gaussian kernel with a full width at half maximum (FWHM) of 6mm. Temporal high-pass filtering utilized a cutoff threshold of 100 seconds.

A two-step co-registration process aligned the functional data first to the participant’s own anatomical image and subsequently to a standardized anatomical template (MNI152). Subjects were defined as having high motion if their mean relative root-mean square displacement exceeded 0.5 mm. For those with excessive motion, displacement plots (mean and relative) were visually examined, and subjects were excluded from further analysis if over 30 consecutive volumes were significantly affected by high motion.

#### First-level analyses and group-mean effects of task

Subject-level analyses were performed using the FEAT module in FSL, employing the general linear model and pre-whitening with FMRIB’s Improved Linear Model (FILM). The design matrix featured distinct explanatory regressors for switch and no-switch trials, each aligned with the cue onset that indicated the trial type. Additionally, six standard head motion parameters were included as nuisance regressors to account for potential confounds caused by motion. Regressors related to the task were convolved with a standard Gamma function (standard deviation = 3s, mean lag = 6s) and modified by adding a temporal derivative and temporal filtering, aligning with the preprocessing steps performed on the data [56]. To model the effects of task switching relative to the no switch component of the time series a switch>no-switch contrast was computed.

Group-level analyses were conducted using FSL’s FLAME-1 module to examine the effects of task switching across participants. Statistical significance for group-level activation was assessed using a whole-brain, cluster-corrected significance threshold (defining threshold Z=2.3, family-wise error corrected at p<0.05)[57].

#### Generation of Regions of Interest

Regions of interest (ROIs) were defined identically to [58]. This was done *a priori* based on an automated meta-analysis of 193 studies identified using neurosynth.org using the term ‘switching’ and a uniformity test (https://neurosynth.org/analyses/terms/switching/). Neurosynth performs automated extraction of activation coordinates from published studies that are associated with specific terms, normalizing these coordinates into a uniform stereotactic space to facilitate meta-analysis. A false-discovery-rate (FDR) correction set at 0.05 is applied to discern differential activation in studies mentioning a term compared to those that do not [59].

For the term ‘switching,’ the resulting activation map was separated into six task-relevant regions, each distinct with no spatial overlap as shown in supplementary figure 2. These regions of interest (ROIs) were converted into binary masks, which were then back-projected into the individual space of each subject. Parameter estimate values were extracted for each region of interest (ROI) mask and for each subject using the featquery module in FSL.

#### Exploratory analyses across different levels of BOLD response

To explore relationships between mitochondrial complex I levels and task switching-related BOLD response across different activation levels, the switch>no switch group level mean parameter estimate map (thresholded at *Z*=2.3, cluster-corrected at p<0.05) was separated into areas that consisted of voxels in different activation ranges. Fslstats was used to define thresholds for four quartiles, Q1 - the bottom 25% of activated voxels, Q2 – voxels in the 25-50% range, Q3, voxels in the 50-75% range and Q4 – top 25% of activated voxels, as well as for the top 1% of activated voxels. These thresholds were applied to create masks for each activation range which were then applied to extract measures of [18F]BCPP-EF VT and switch>no switch PE for each subject using the same approach as described above.

### Statistical analyses

#### Relationship between [18F]BCPP-EF VT and switching-evoked neural activation

Our primary hypothesis investigates the statistical association between brain activity during task switching, as detected by the fMRI BOLD response, and the levels of mitochondrial complex I, measured using [18F]BCPP-EF PET. Given that both fMRI and PET data include measures for multiple ROIs for each participant, we applied a Partial Least Squares Canonical Analysis (PLS-CA) to analyse the multivariate relationships across the six ROIs across the two neuroimaging modalities, as previously implemented in [58]. The PLS-CA approach is particularly well-suited for neuroimaging data, which typically includes many inter-correlated variables [60]. PLS-CA was implemented using the ‘PLSCanonical’ function from the ‘sklearn.cross_decomposition’ module in Python [61].

In summary, the PLS-CA approach computes matrices for each modality ([n_participants, n_ROIs]) and identifies linear projections of the fMRI and PET data where for each new multidimensional direction *i* the projected fMRI and PET datasets (referred to as fMRI and PET ‘*scores_i_*’) exhibit maximal covariance. This is done under the constraint that scores from different projections remain orthogonal to each other. We used a two-component model (i.e., identifying solutions which yielded projections of fMRI and PET datasets onto a two-dimensional subspace) allowing us to investigate the first and second most correlated latent structures between the fMRI and PET data. The two-component solution allowed balancing model simplicity and the ability to explain the shared variance between the PET and fMRI datasets.

To assess the statistical significance of the identified relationships between PET and fMRI data (canonical correlations), we utilized a permutation testing method, described in [58]. This approach robustly calculates p-values by comparing observed canonical correlations with a null distribution of correlations derived from randomly permuting the rows of the fMRI [n_participants, n_ROI] matrix several times, while keeping the rows of the PET [n_participants, n_ROI] matrix constant. For this study, we executed 1000 permutations and set a significance threshold at p<0.05. In addition to reporting canonical correlations, we also report canonical weights for each latent component. These weights are a [1, number_ROI] vector for both fMRI and PET and signify the contribution of each original feature (from each ROI) to the canonical variates, allowing us to understand which brain regions primarily drive the shared information between the PET and fMRI data.

To investigate the nature of the relationships at the individual ROI level, we carried out exploratory analyses calculating Pearson’s correlation coefficients to assess relationships between [18F]BCPP-EF PET and task switching fMRI and performance measures. Similarly, to address our exploratory hypotheses of whether the strength of relationships between [18F]BCPP-EF VT and the task switching neural response may be related to the level of neural activity, we used switch>no switch PE values and [18F]BCPP-EF VT extracted from voxels at different activation ranges (described above) and carried out Pearson’s correlation to assess the relationship between these measures. *P* values were uncorrected for exploratory analyses and results are discussed as significant if p<0.05.

#### Relationship between [18F]BCPP-EF VT and switching task performance

To explore the associations between [18F]BCPP-EF PET data and task performance metrics, we employed Partial Least Squares Regression (PLS-R), as the method is suitable for neuroimaging data where predictors exhibit multicollinearity (e.g [18F]BCPP-EF PET data). Similarly to the PLS-CA, in this approach, both PET and task performance data are projected onto a new set of orthogonal components designed to maximize the covariance between them. This technique helps identify latent structures that most significantly relate mitochondrial complex I levels to task switching performance. We focused on two specific performance measures, switch cost and switching accuracy, conducting separate PLS-R analyses for each, each utilizing two components.

The predictive accuracy of our models was evaluated using the R^2^ statistic and the root mean square error (RMSE). Additionally, we integrated a permutation test (1000 permutations) to provide a non-parametric evaluation of the R^2^ values of our models. We additionally conducted exploratory univariate analyses, specifically we calculated Pearson’s correlation coefficients to assess the associations between PET data and task performance, applying a significance threshold of p<0.05.

## Results

### 3.1 Demographic details and effects of task

A total of 25 subjects were recruited and completed the study. 23 participants were included in the final analyses, as one was excluded due to non-performance on the task and one due to high motion (see methods for details). Behavioural data for one subject were not available due to an MRI controller issue. Full sample details are presented in supplementary table 1.

Mean whole-brain [18F]BCPP-EF VT is shown in figure 1A for reference, indicating mitochondrial complex I levels across the brain, with injected dose information provided in supplementary table 2. As shown in figure 1B, the task engaged relevant regions including regions preselected as ROIs (supplementary figure 2); these were the dorsolateral prefrontal cortex, insula, parietal cortex, posterior frontal cortex, anterior cingulate cortex, thalamus, and putamen. All preselected ROIs had positive mean parameter estimate values (supplementary figure 3).

**Figure 1.**
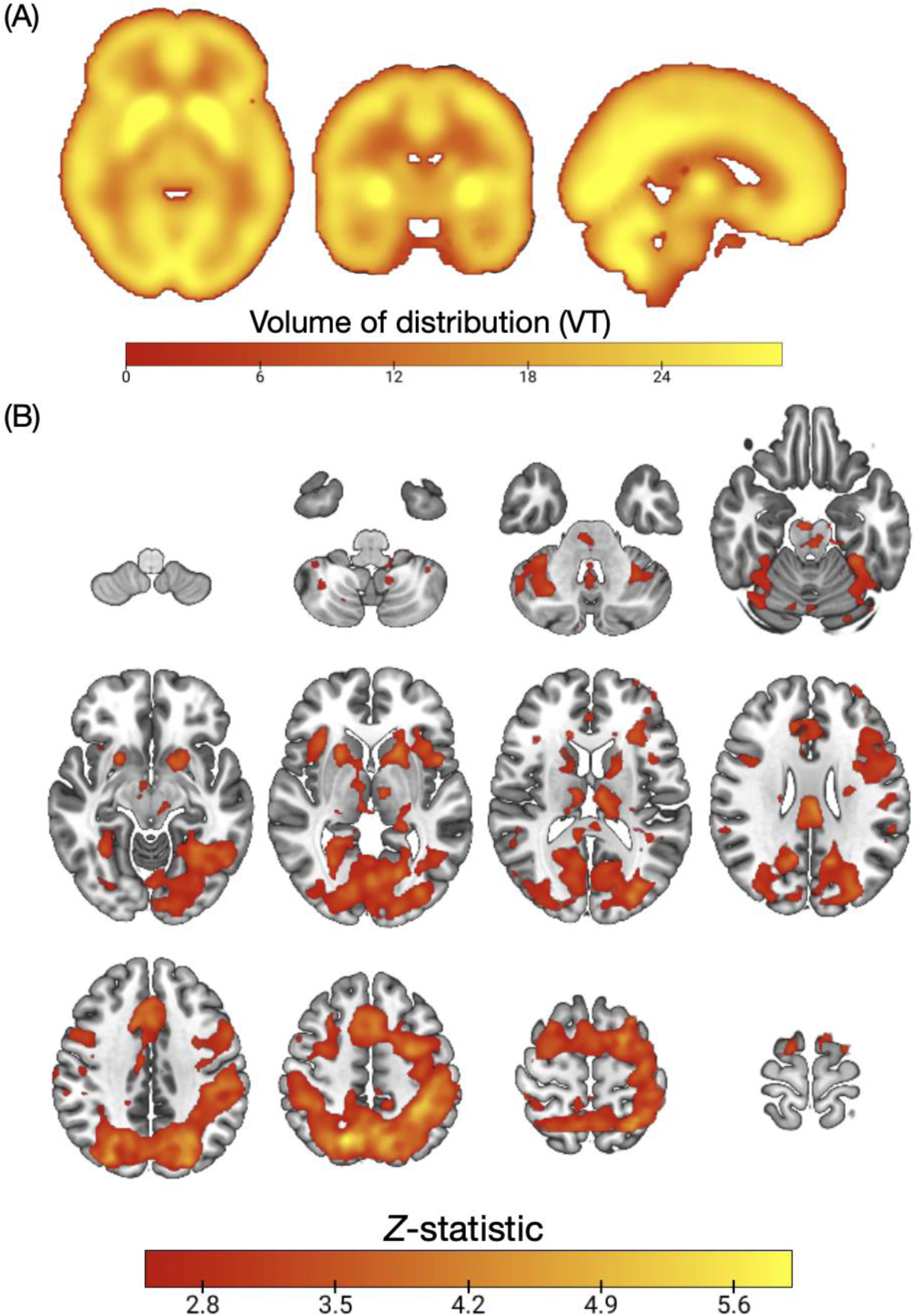
A: Mean [18F]BCPP-EF mean volume of distribution (VT) (*n*=23). **1B: Mean activation during task switching (left, n=23) subjects:** Z (Gaussianised T/F) statistical images representative of significant activation during the Switch>noSwitch condition of the switching task. Images were thresholded using clusters determined by Z > 2.3 and a (corrected) cluster significance threshold of p = 0.05, Axial slices shown in MNI152 are: -60 -46 -34 -22; -10 2 14 26; 38 50 62 74.

### 3.2 Mitochondrial complex I and task switching BOLD response

As shown in figure 2, we found that in the first canonical component, which captures the greatest amount of covariance in our data, there was a statistically significant relationship between [18F]BCPP-EF VT and switch>no-switch (3A, Cov=2.73, *r*=0.51, *p*=0.030). As shown in supplementary table 3, fMRI data from the different ROIs contributed similarly to the first canonical variate (weights ranging between 0.341-0.475), while for the PET data, the thalamus-putamen and insula ROI PET data contributed most to the first canonical variate (weights 0.759 and 0.519, respectively). The correlation between PET and fMRI second canonical variates was not significant (3B, Cov=0.12, *r*=0.57, *p*=0.214).

**Figure 2:**
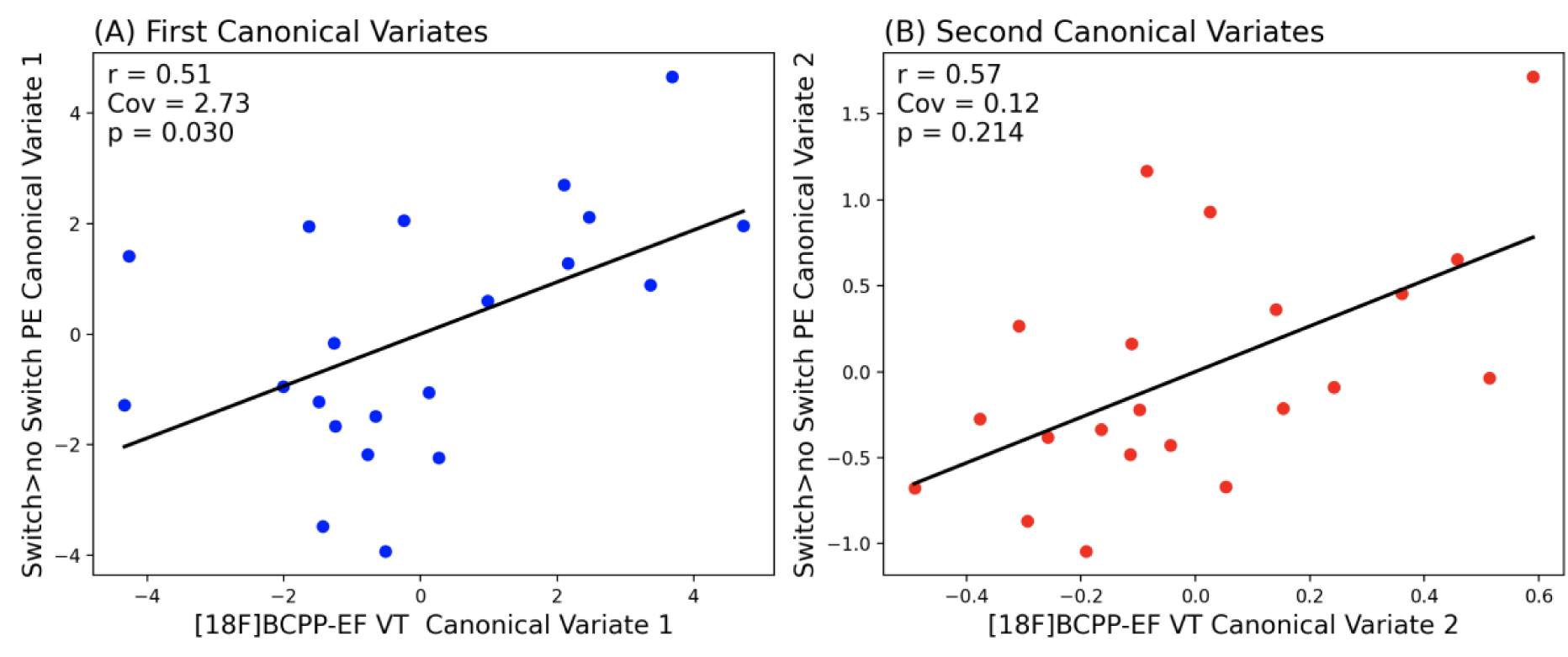
Partial Least Squares Canonical Analysis (PLS-CA) showing a positive correlation between [18F]BCPP-EF volume of distribution (VT) and task switching related neural activity (Switch>noSwitch parameter estimates) in the first canonical component (A) and no significant relationship in the second canonical component (B). Cov – covariance captured by each canonical correlation, significance threshold p<0.05, PE – parameter estimates.

**Figure 3:**
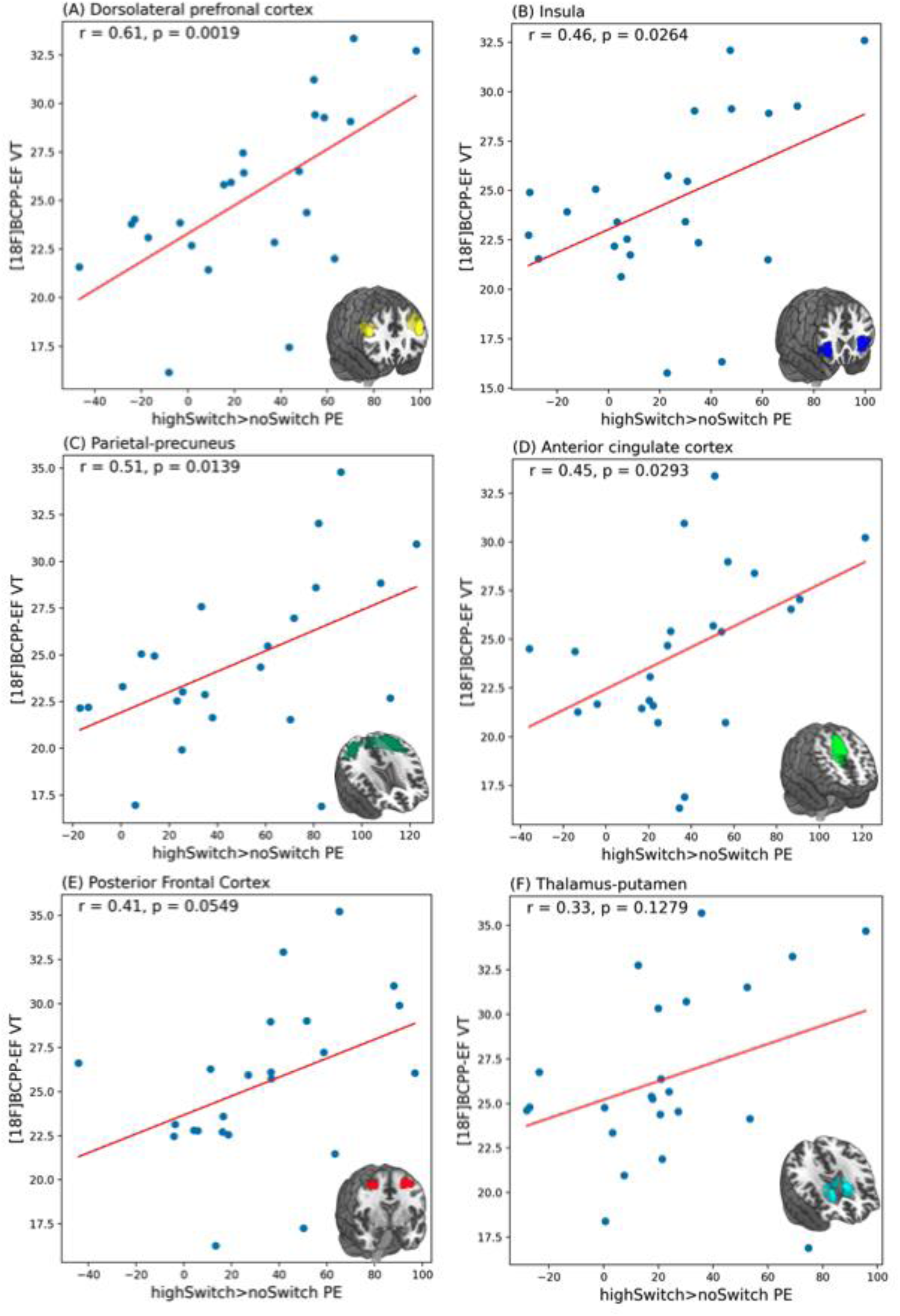
Scatter plots and Pearson’s correlation results showing significant positive relationship between [18F]BCPP-EF VT and switch>no-switch PE in the dorsolateral prefrontal cortex (**A**), insula (**C**), parietal-precuneus (**D**), anterior cingulate cortex (**E**), and lack of significant relationships between [18F]BCPP-EF VT and switch>no-switch PE in the posterior frontal cortex ROI (**B**) and thalamus-putamen regions of interest.

Exploratory Pearson’s correlations showed positive correlations between [18F]BCPP-EF VT and switch>no-switch PE in the DLPFC (*r*=0.61, *p*=0.0019, 3A), Insula (*r*=0.46, *p*=0.0264, 3B), parietal-precuneus (*r*=0.51, *p*=0.0139, 3C) and anterior cingulate cortex (*r*=0.45, *p*=0.0293, 3D) ROIs. These relationships did not reach significance in the posterior frontal cortex (*r*=0.41, *p*=0.0549, 3E) and thalamus-putamen (*r*=0.33, *p*=0.1279, 3F) ROIs.

Having established that intersubject variability in mitochondrial complex I levels is related to task switching BOLD signal, we proceeded to test for a “dose” relationship with the strength of the BOLD response across the brain. This approach goes beyond the PLS-CA approach as it captures how relationships between [18F]BCPP-EF VT and switch>no-switch PE change across the brain depending on the level of activation of the voxels. The switch>no switch group level mean parameter estimate map (thresholded at Z>2.3, cluster p<0.05, shown in figure 1) was separated into areas that consisted of voxels in different activation levels: four quartiles, Q1 - the bottom 25% of activated voxels, Q2 – voxels in the 25-50% range, Q3, voxels in the 50-75% range and Q4 – top 25% of activated voxels, and top 1% of activated voxels. Masks for these regions are shown in supplementary figure 5. [18F]BCPP-EF VT and switch>no switch PE for each subject were extracted from voxels in each of these activation ranges. We found that in the two lowest quartiles of activated voxels, relationships between [18F]BCPP-EF VT and switch>no switch PE did not reach significance (Q1 *r*=0.37, *p*=0.0864, 6A; Q2 *r*=0.40, *p*=0.0598, 6B). But in the top two quartiles and top 1% of activated voxels [18F]BCPP-EF VT and switch>no switch PE were positively correlated (Q3 *r*=0.42, *p*=0.0439, 6C; Q4 *r*=0.49, *p*=0.0195, 6D; top 1% *r*=0.54, *p*=0.0080, 6E). Pearson’s *r* coefficients increased for each consecutive correlation from lower to higher activation ranges, while *p* values decreased in a similar manner (figure 4).

**Figure 4:**
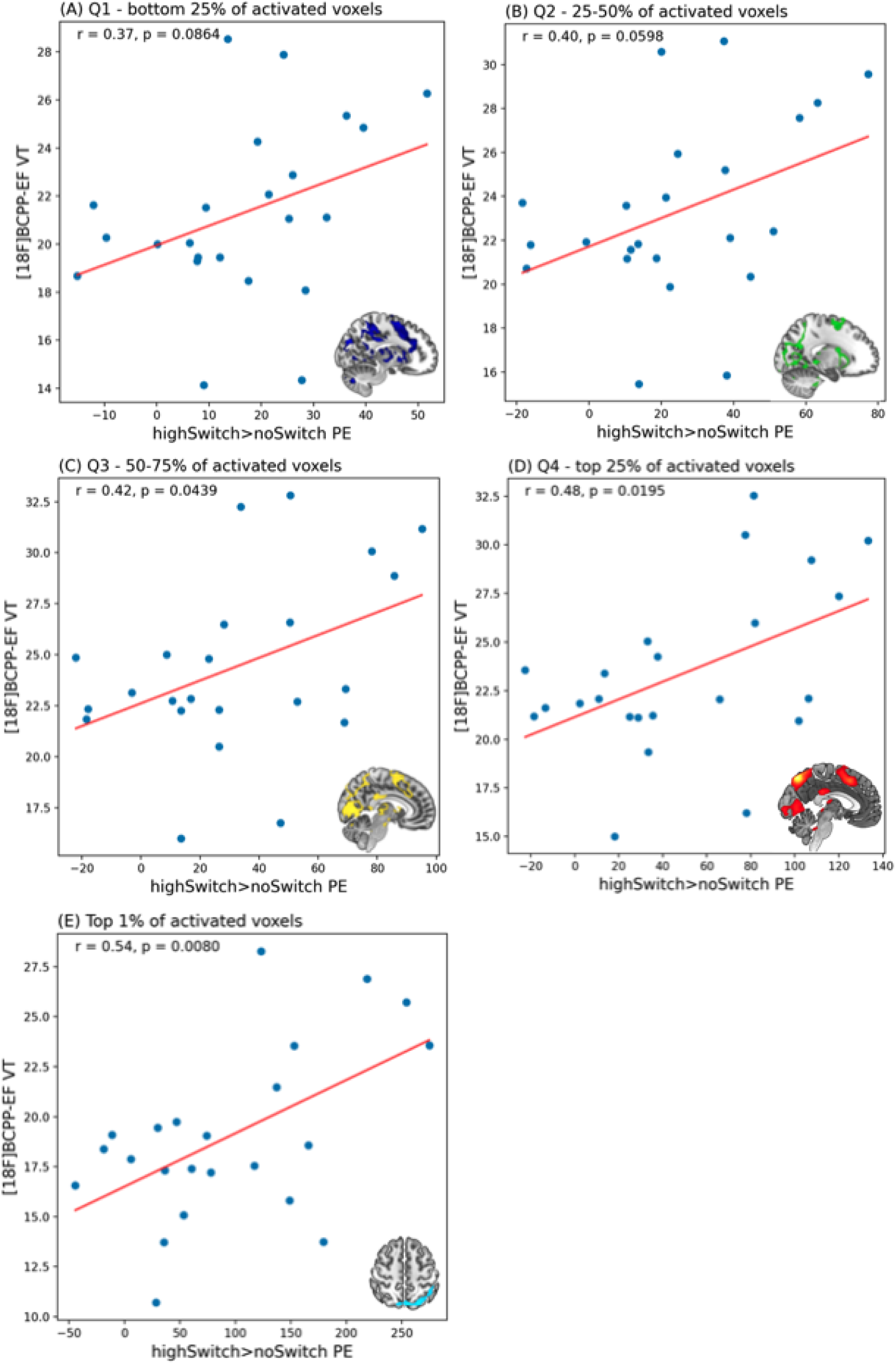
Scatter plots showing [18F]BCPP-EF VT and switch>no switch PE in different activation ranges. (A) Lower quartile, bottom 25% (Q1), (B) 25-50% (Q2), (C) 50-75% (Q3), (D)top 25 % quartile (Q4), (E) Top 1%. [18F]BCPP-EF VT and switch>no switch PE are extracted from identical regions for each subject indicated on each plot and in supplementary figure 4.

### 3.3 Mitochondrial complex I and task switching performance

As shown in figure 5, [18F]BCPP-EF VT values from the six task switching regions of interest were significantly predictive of both task switching accuracy (PLS-regression, R^2^=0.48, RMSE=0.14, *p*=0.011, 6A), and of switch cost (PLS-regression, R^2^=0.38, RMSE=0.07, *p*=0.048, 6B). In the first component of the PLS-R for accuracy (covariance captured 0.043) the PET data for the thalamus-putamen had the highest weight (0.930, supplementary table 5), while for the switch-cost PLS-R the anterior cingulate [18F]BCPP-EF VT has the highest weight (0.723 supplementary table 4). Exploratory analyses showed no significant positive or negative correlations between [18F]BCPP-EF VT and switch cost or switching accuracy in individual ROIs (supplementary table 6).

**Figure 5:**
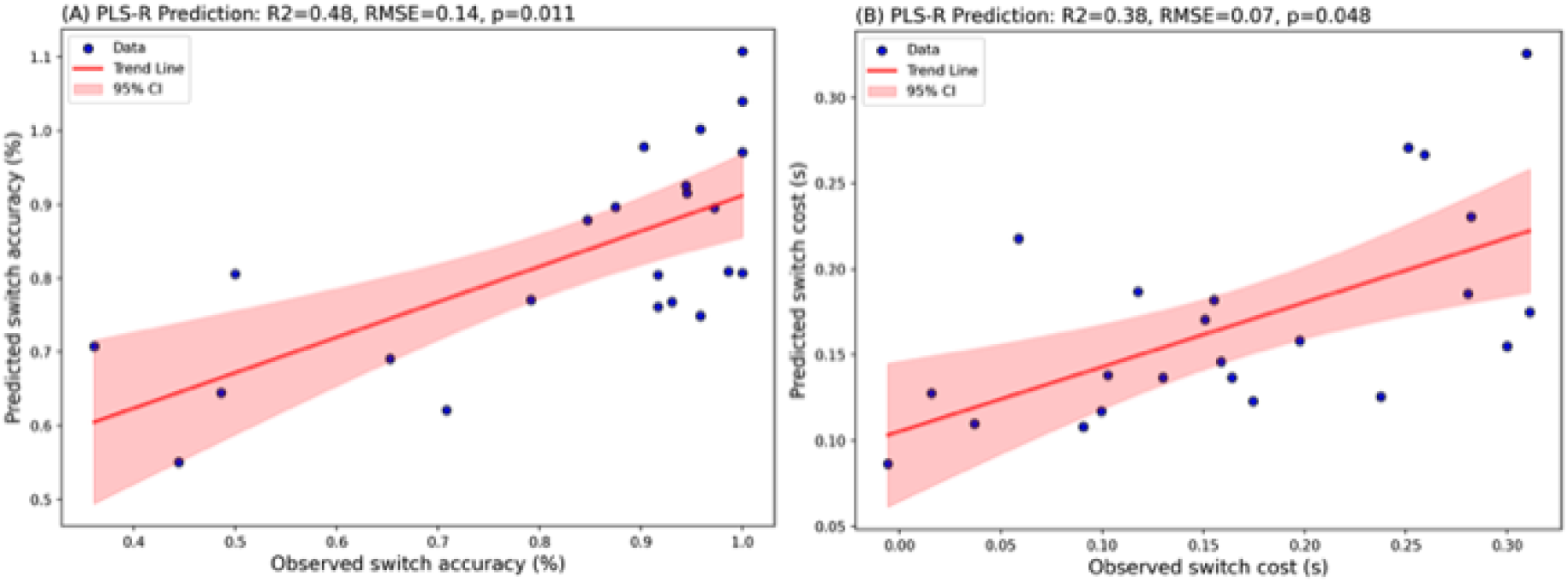
Two-component partial-least-squares regression results showing (A) switching accuracy and (B) switch cost (s) are significantly predicted by [18F]BCPP-EF VT values across six task-switching relevant regions of interest. Figure shows predicted values plotted against observed values, trend-lines are in red and 95% confidence intervals are shaded red.

## Discussion

In this study we have shown for the first time that, in the healthy brain, that task switching BOLD signal was positively related to mitochondrial complex I levels in brain regions related to switching. Importantly this is the first time that a direct relationship between the density of a mitochondrial respiratory chain component and BOLD signal has been reported in humans. The BOLD signal captures the haemodynamic and vascular response to increased oxygen demand of activated neurones, driven by the energy demand of synaptic firing [21–24]. Given that electron microscopy studies show synapses are densely packed with mitochondria [62], our findings are in agreement with what is known about the biological basis of BOLD. They add to this body of literature by suggesting individuals with higher mitochondrial complex I levels are able to produce a stronger BOLD response.

Exploratory analyses showed a significant positive correlation between BOLD signal and mitochondrial complex I levels in the DLPFC, ACC, parietal-precuneus and insula and a trend towards a positive correlation in posterior frontal regions (*p*=0.05). Importantly, the strongest positive association between mitochondrial complex I levels and BOLD signal was in the DLPFC. Previous work has shown that the frontoparietal network, particularly the DLPFC is closely related to task switching [40, 63–66], specifically in task maintenance and top-down control [67]. Moreover, transcranial direct current stimulation studies have also shown that stimulation of the DLPFC can improve task switching performance [68] and that switch cost correlated with DLPFC hypometabolism in Parkinson’s disease patients [69]. Our interpretation for these findings is that the DLPFC of individuals with higher mitochondrial complex I levels may be metabolically more efficient and that this efficiency permits greater or more sustained neuronal activity, resulting in higher BOLD signals during task switching. In line with this, we found that mitochondrial complex I levels were significantly predictive of both switch cost and switching accuracy when evaluating data from all six ROIs together. While we did not see a relationship between mitochondrial complex I levels and switch cost in specific ROIs, one reason for this might be that in healthy humans there is little between subject variability in task performance measures, and relationships in individual regions may become more pronounced in cognitively impaired populations.

In further exploratory analyses, we found that the relationship between mitochondrial complex I and switching-related BOLD was strongest in voxels where BOLD signal was highest. Specifically, in the lowest 50% of activated voxels (two lowest quartiles) the correlation did not reach significance, but for each consecutive activation range (50-75%, top 25%, top 1%), the Pearson’s correlation *r* coefficients increased, and *p* values decreased indicating a stronger relationship. One possible interpretation of these findings is that the oxygen demand is related to increased synaptic activity and therefore there is a stronger dependency on total mitochondrial complex I availability in regions most engaged by the task. On the other hand, a key consideration is that going up the activation ranges for fMRI captures an increasing signal-to-noise ratio. For example, earlier fMRI studies of cognitive function have shown that evaluating data from top 20% activated voxels produces more robust and reproducible estimates of task associated activity [70]. This could therefore suggest that at lower activation ranges, the relationship is not significant due to increased noise in the data.

Taken together these findings extend the body of research investigating the biological underpinnings of BOLD signal [21–24, 71], by suggesting individual BOLD response to task switching may be a function of mitochondrial complex I availability, especially at higher activation ranges. This proposes that mitochondrial complex I availability may be a key constraint on BOLD signal increases during executive function, and replicating these results in other tasks could confirm their generalizability beyond task switching. Additionally, previous work also shows that task-related increases in BOLD are typically signal changes below 1-5% of the baseline signal. Thus, while we link interindividual differences in mitochondrial complex I levels with differences in task-related BOLD signal change, a key question that arises is whether mitochondrial complex I levels may also explain differences in baseline BOLD signal. Thus, a key next step would be to evaluate these relationships during resting conditions.

### Strengths and limitations

Firstly, [18F]BCPP-EF binds to mitochondrial complex I with high affinity (K_I_=2.3), and in competition studies is almost completely displaced by rotenone, which is known to be a highly specific inhibitor of mitochondrial complex I function [44, 49, 72, 73]. This suggests that non-specific binding for [^18^F]BCPP-EF in the mammalian brain is low and that signal measured in this study is specific to mitochondrial complex I. Nonetheless, given that it is a novel PET probe, additional validation will be required, although available evidence suggests it has good test-retest reliability, providing additional confidence in our results [43].

While the radiotracer is specific to mitochondrial complex I, it important to note that other biological parameters may impact mitochondrial complex I levels for an individual. For example, the positive relationship between BOLD signal and mitochondrial complex I, could also partially capture individual differences in mitochondrial number or in synaptic density. Joint [18F]BCPP-EF PET fMRI studies in clinical populations with known mitochondrial complex I pathology would support the specificity of the relationships we report.

One other limitation is the sample size of this study. While this has sufficient power to detect moderate or larger associations it lacks power to detect weak associations, which may account for our finding that mitochondrial complex I Ievels were not related to switch cost and switch accuracy in individual regions. However, the observed *r* values were low (<0.2) and, given the small amount of variance that could be explained, it is unlikely such weak relationships would be meaningful in terms of understanding mechanisms. Finally, in addition to this, given that our sample included few female participants (∼10%), it is important to replicating these findings in a larger mixed cohort to see if these findings may generalise.

### Implications

Our findings suggest that BOLD signal change associated with engaging executive function is related to mitochondrial complex I levels, especially in regions that are highly activated by the task. Schizophrenia, Parkinson’s disease, Alzheimer’s disease, and other conditions have been found to be associated with mitochondrial complex I dysfunction [5–9]. Interestingly, they are also associated with aberrant BOLD responses during executive function tasks [16–20]. For example, individuals with schizophrenia have been shown to have a lower DLPFC BOLD response during executive function tasks upon meta-analysis [74]. Similarly, patients with Parkinson’s disease show lower activation of the inferior frontal gyrus pars opercularis during cognitive tasks, upon systematic review [75]. Given that these conditions are also associated with alterations in mitochondrial complex I, the findings of our study propose a possible biological mechanism that could explain aberrant BOLD signal in these conditions. Thus, testing mitochondrial complex I–BOLD signal relationships in these clinical cohorts would be a key next step to understand if mitochondrial complex I dysfunction might be driving the aberrant BOLD responses in these populations.

### Conclusions

Together our findings show that levels of mitochondrial complex I are related to the strength of an individual’s blood oxygen level dependent response during an executive function task. Given this, it is crucial to test whether that mitochondrial complex I pathology in conditions such as Parkinson’s disease, Alzheimer’s disease and schizophrenia may be contributing to the aberrant blood oxygen level dependent response reported during an cognitive tasks in these conditions.

## Author contributions

Study concept and design: ES, TW, OH

Data collection: ES, TW, ECO, GR, AW, AM, TRM, SN, ER, MW, OH

Acquisition, analysis, or interpretation of data: ES, TW, ECO, GR, AW, AM, TRM, SN, ER, MW, OH – all authors

Drafting of the manuscript: ES, TW, ECO, GR, AW, AM, TRM, SN, ER, MW, OH – all authors

Critical revision of the manuscript for important intellectual content: ES, TW, ECO, GR, AW, AM, TRM, SN, ER, MW, OH – all authors

Statistical analysis: ES, TW

Study supervision: OH

## Data Availability statement

The datasets used and/or analysed during the current study are available from the corresponding author on reasonable request.

## Acknowledgements/disclosures

Not applicable

