## Supplement for "The relationship between brain activation and mitochondrial complex I protein levels during cognitive function in healthy humans: an [18F]BCPP-EF PET and functional MRI study of task switching"


**This PDF file includes:**

Supporting text
Figures S1 to S4
Tables S1 to S6
SI References

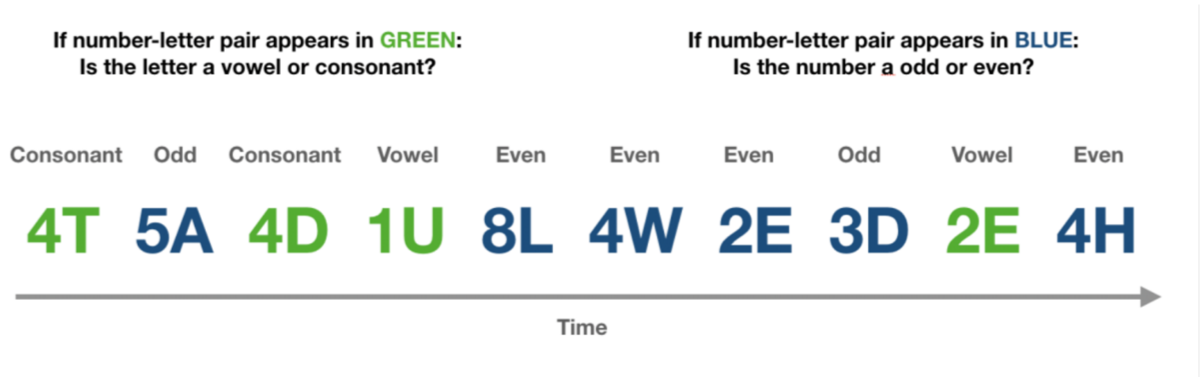


**Supplementary figure 1: Task switching executive function task.** Pairs of numbers and letters appeared on the screen in blue or green. If the number-letter pair appeared in green, participants were required to focus on the letter and respond if the letter is a vowel or consonant using an MRI-compatible response box, as quickly as possible. If the number-letter pair appeared in blue, participants were required to focus on the number and response if it is odd or even. Cue colours/tasks switched pseudo-randomly.

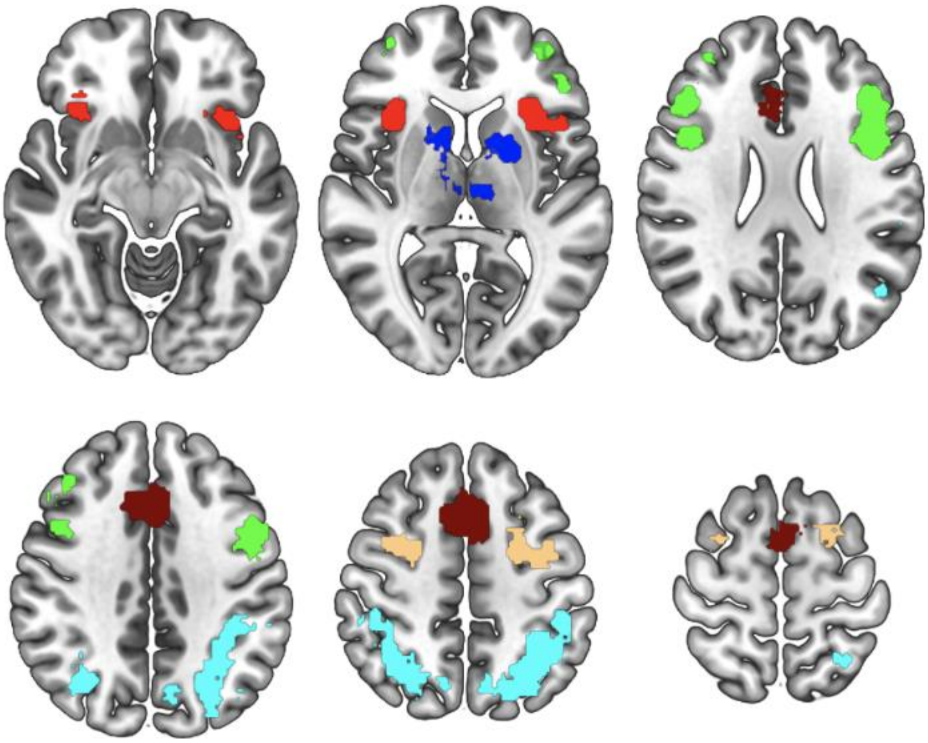

**Supplementary figure 2: Task switching a priori selected ROIs:** Dorsolateral prefrontal cortex (green), insula (red), parietal cortex (light blue), posterior frontal cortex (beige), anterior cingulate cortex (maroon), thalamus-putamen (dark blue). Shown in MNI152 space, slices are: A -10 7 25; 38 50 62

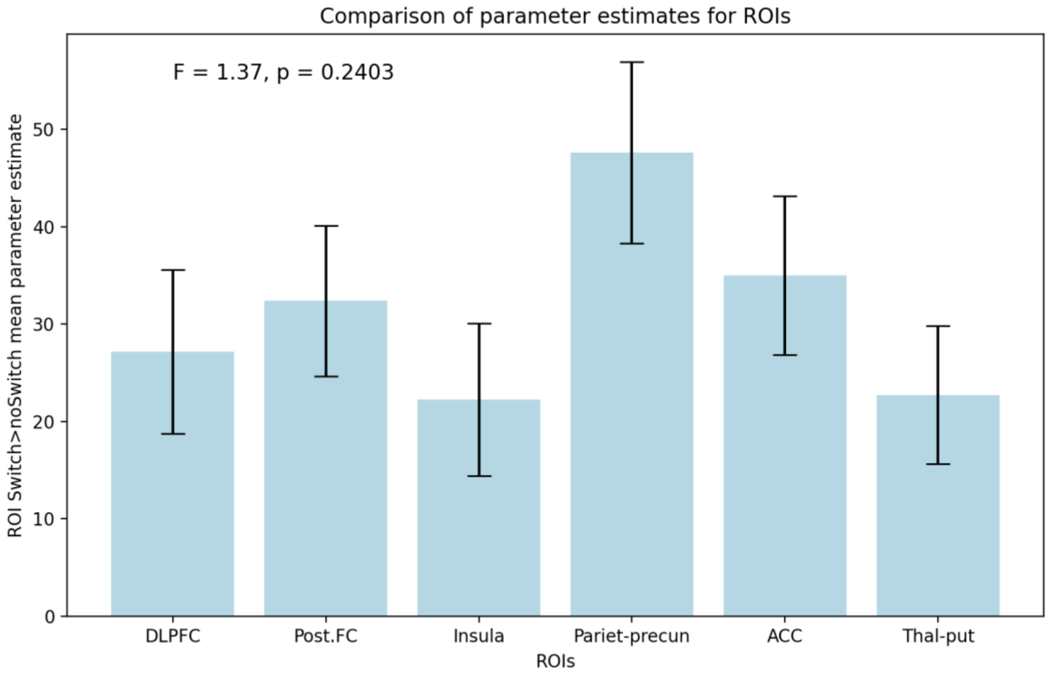

**Supplementary figure 3:** Mean parameter estimates for each region of interest for switch>no-switch contrast. All regions have positive paramater estimates that are comparable across regions (one-way ANOVA, F=1.37, p=0.2403). Error bars = standard error of the mean.

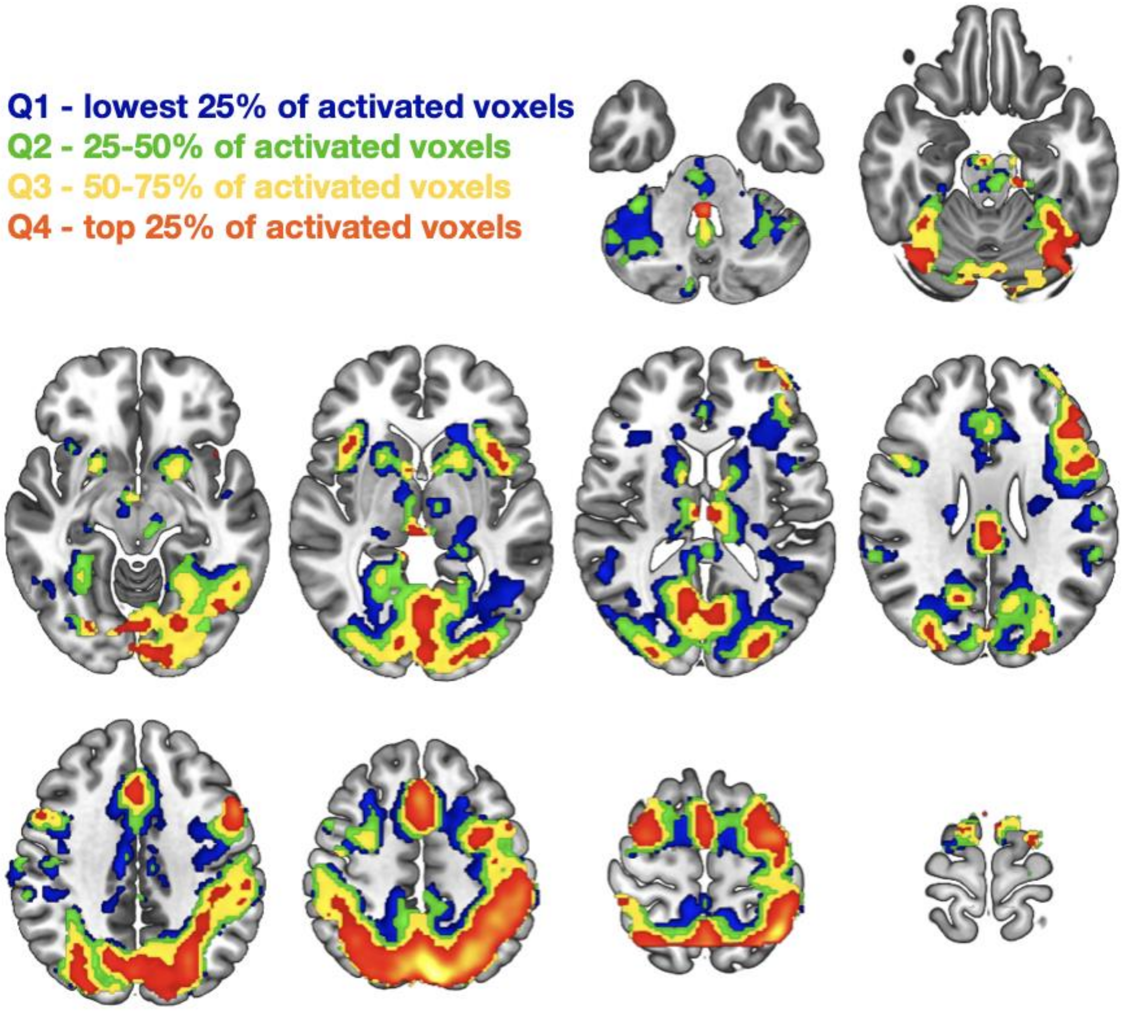


**Suplementary figure 4:** Masks corresponding to task switching-related BOLD signal split into quartiles. Lower quartile (Q1) in blue, 25-50% (Q2) in green, 50-75% (Q3) in yellow, top quartile (Q4) in orange-red

**Supplementary table 1:** Demographic details

| | Mean ± Standard Deviation |
|---|---|
| Sample Size | *n*=23 |
| Male/Female | 21/2 |
| Age (years) | 35.17 ± 12.21 |
| Weight (kg) | 83.01 ± 15.27 |
| BMI (kg/m$^2$) | 26.28 ± 4.55 |
| Injected radioactivity (MBq) | 91.49 ± 3.27 |
| Plasma fraction | 0.24 ± 0.02 |

**Supplementary table 2:** Details of [18F]BCPP-EF injected activity, injected mass and specific activity for each subject

| Subject id | Age | Sex | Weight | Injected Activity (MBq) | Injected mass (µm) | Specific activity (GBq/µmol) |
|---|---|---|---|---|---|---|
| 901 | 53 | M | 85.9 | 97.4 | 0.12 | 334.7 |
| 903 | 25 | M | 69.3 | 91.3 | 0.05 | 718.0 |
| 907 | 27 | M | 78.9 | 92.3 | 0.04 | 903.2 |
| 908 | 23 | M | 69.3 | 90.4 | 0.15 | 248.0 |
| 911 | 59 | M | 78.1 | 86.8 | 0.1 | 339.6 |
| 912 | 52 | M | 82 | 91.5 | 0.07 | 527.7 |
| 913 | 42 | M | 93.6 | 88.5 | 0.11 | 335.0 |
| 915 | 39 | M | 94.1 | 86.6 | 0.06 | 600.0 |
| 917 | 39 | M | 113.3 | 90.6 | 0.06 | 639.5 |
| 919 | 49 | M | 93.2 | 96.2 | 0.23 | 170.3 |
| 921 | 34 | M | 112 | 91.1 | 0.14 | 264.3 |
| 922 | 48 | M | 73 | 93.5 | 0.14 | 276.2 |
| 923 | 50 | M | 93.8 | 96.6 | 0.08 | 478.0 |
| 927 | 25 | M | 106.3 | 93.5 | 0.15 | 253.4 |
| 929 | 30 | M | 70.2 | 83.6 | 0.08 | 426.5 |
| 930 | 38 | M | 98.8 | 90.2 | 0.16 | 227.1 |
| 931 | 40 | M | 77.5 | 88.7 | 0.16 | 219.6 |
| 935 | 21 | F | 63.7 | 92.6 | 0.12 | 319.8 |
| 943 | 28 | M | 69.7 | 94.1 | 0.05 | 804.9 |
| 944 | 22 | M | 62.7 | 90.8 | 0.05 | 670.2 |
| 950 | 22 | M | 69.8 | 92.2 | 0.11 | 325.2 |

| | | | | | | |
|---|---|---|---|---|---|---|
| 956 | 20 | F | 67.1 | 91.9 | 0.18 | 203.5 |
| 959 | 23 | M | 86.9 | 94.0 | 0.25 | 152.6 |

**Supplementary Table 3:** Canonical weights for [18F]BCPP-EF PET and task switching fMRI data included in the PLS-CA analysis

| | | Insula | DLPFC | Parietal-precuneus | Post. Front. Ctx | ACC | Thalamus-putamen |
|---|---|---|---|---|---|---|---|
| First component | PET | 0.387 | 0.389 | 0.428 | 0.475 | 0.416 | 0.341 |
| | fMRI | 0.519 | 0.216 | 0.154 | 0.072 | 0.281 | 0.759 |
| Second component | PET | 0.041 | 0.050 | 0.239 | -0.500 | -0.369 | 0.744 |
| | fMRI | -0.291 | 0.385 | 0.657 | 0.533 | 0.166 | -0.156 |

**Supplementary Table 4:** Canonical weights for [18F]BCPP-EF PET included in the PLS-R analysis testing if [18F]BCPP-EF VT is predictive of switch cost

| | | Insula | DLPFC | Parietal-precuneus | Post. Front. Ctx | ACC | Thalamus-putamen |
|---|---|---|---|---|---|---|---|
| switch cost | First component | 0.119 | 0.377 | 0.151 | 0.502 | 0.724 | 0.213 |
| | Second component | 0.585 | 0.173 | 0.539 | -0.034 | -0.398 | 0.420 |

**Supplementary Table 5:** Canonical weights for [18F]BCPP-EF PET included in the PLS-R analysis testing if [18F]BCPP-EF VT is predictive of switch accuracy

| | | Insula | DLPFC | Parietal-precuneus | Post. Front. Ctx | ACC | Thalamus-putamen |
|---|---|---|---|---|---|---|---|
| switch accuracy | First component | 0.106 | -0.229 | 0.184 | -0.189 | 0.002 | 0.931 |
| | Second component | 0.405 | 0.521 | 0.370 | 0.486 | 0.423 | 0.107 |

**Supplementary Table 6:** Exploratory Pearson's correlations between [18F]BCPP-EF VT in individual ROIs and task switching performance.

| Variable | Region of interest | Pearson's r | p-value |
|---|---|---|---|
| Switch cost | Dorsolateral prefrontal cortex | 0.129 | 0.558 |
| | Posterior frontal cortex | 0.171 | 0.435 |
| | Insula | 0.040 | 0.855 |
| | Parietal-precuneus | 0.051 | 0.816 |
| | Anterior cingulate cortex | 0.247 | 0.256 |
| | Thalamus-putamen | 0.072 | 0.742 |
| Switching accuracy | Dorsolateral prefrontal cortex | -0.048 | 0.827 |
| | Posterior frontal cortex | -0.040 | 0.856 |
| | Insula | 0.022 | 0.919 |
| | Parietal-precuneus | 0.039 | 0.861 |
| | Anterior cingulate cortex | 0.000 | 0.998 |
| | Thalamus-putamen | 0.196 | 0.369 |

**Calculation of supraphysiological threshold for [$^{18}$F]BCPP-EF**

Outlier voxels are an issue when applying the MA1 model in a voxelwise manner. To calculate a threshold for removing outlier voxels we used mean volume of distribution (VT) values previously published for [$^{18}$F]BCPP-EF for a range of brain regions [1] (see [2] for additional details), followed by calculating the standard deviation and calculating a mean±3SD VT value for each region, which was assumed as a supraphysiological threshold. The highest uptake for the tracer is in the putamen, where the mean±3SD VT value was 53.38. This was rounded to 55 and used as a threshold for our data to remove outlier voxels. No excluded voxels overlapped with the preselected ROIs used in this study.
